# Mice with High FGF21 Serum Levels Had a Reduced Preference for Morphine and an Attenuated Development of Acute Antinociceptive Tolerance and Physical Dependence

**DOI:** 10.1101/2021.07.30.453845

**Authors:** Louben Dorval, Brian I. Knapp, Olufolake A. Majekodunmi, Sophia Eliseeva, Jean M. Bidlack

## Abstract

Because of increased opioid misuse, there is a need to identify new targets for minimizing opioid tolerance, and physical and psychological dependence. Previous studies showed that fibroblast growth factor 21 (FGF21) decreased alcohol and sweet preference in mice. In this study, FGF21-transgenic (FGF21-Tg) mice, expressing high FGF21 serum levels, and wildtype (WT) C57BL/6J littermates were treated with morphine and saline to determine if differences exist in their physiological and behavioral responses to opioids. FGF21-Tg mice displayed reduced preference for morphine in the conditioned place preference assay compared to WT littermates. Similarly, FGF21-Tg mice had an attenuation of the magnitude and rate of acute morphine antinociceptive tolerance development, and acute and chronic morphine physical dependence, but exhibited no change in chronic morphine antinociceptive tolerance. The ED50 values for morphine-induced antinociception in the 55°C hot plate and the 55°C warm-water tail withdrawal assays were similar in both strains of mice. Likewise, FGF21-Tg and WT littermates had comparable responses to morphine-induced respiratory depression. Overall, FGF21-Tg mice had a decrease in the development of acute analgesic tolerance, and the development of physical dependence, and morphine preference. FGF21 and its receptor have therapeutic potential for reducing opioid withdrawal symptoms and craving, and augmenting opioid therapeutics for acute pain patients to minimize tolerance development.

## 1. Introduction

Opioid misuse continues to be a serious problem affecting health, social, and economic welfare nationally and globally. Adverse effects of opioid misuse include analgesic tolerance development, physical and psychological dependence, respiratory depression, withdrawal symptoms, and relapse. Acute pain patients suffer from the development of tolerance, reducing the efficacy of opioid medications. Similarly, some patients continue to take opioids to prevent withdrawal symptoms, a sign of physical dependence. The psychological dependence of craving opioids is associated with opioid misuse. Increases in dopamine levels in certain brain regions mediate the reinforcing effects of opioids (Cao et al., 2021; Nisell et al., 1994; Wiss et al., 2018). In addition to opioids, sucrose, saccharin and alcohol, as well as other drugs of abuse, increase dopamine release in the nucleus accumbens (NAc) and this increase in dopamine levels has been correlated to development of preference in mice (Ma and Zhu, 2014; Yoshimoto et al., 1992). Novel therapeutic targets are needed to attenuate acute opioid tolerance, and physical and psychological dependence, while not affecting opioid-induced analgesia.

Fibroblast growth factor 21 (FGF21) is a 20 kDa protein expressed in liver, brown adipose tissue, glia, and neurons (Potthoff et al., 2012). Unlike most fibroblast growth factors (FGFs), FGF21 is released into the bloodstream and can act throughout the body (Potthoff et al., 2012). FGF21 administration in obese diabetic mice ameliorated hyperglycemia and lowered elevated triglycerides (Coskun et al., 2008). Moreover, FGF21 administered to obese diabetic monkeys improved glucose and circulating lipids levels (Kharitonenkov et al., 2007). FGF21 analogues are now in clinical trials to treat type 2 diabetes, obesity, and nonalcoholic steatohepatitis (Geng et al., 2020). PF-05231023, and Efruxifermin (AKR-001) produced a decrease in triglycerides, total cholesterol, low-density lipoprotein cholesterol and an increase in high-density lipoprotein cholesterol (Dong et al., 2015; Gaich et al., 2013; Stanislaus et al., 2017). Importantly, FGF21 crosses the blood-brain barrier and acts centrally to induce the sympathetic nervous system (Hsuchou et al., 2007; Owen et al., 2014) by binding to FGF receptors in complex with the obligate co-receptor β-Klotho (KLB) (Ding et al., 2012; Owen et al., 2015). In mice, elevated FGF21 levels and the FGF21 analogue PF-05231023 have been shown to attenuate sweet and alcohol preference (Talukdar et al., 2016).

In addition, mouse models have been created to better understand FGF21 signaling pathway. In this study, we used the FGF21 transgenic (FGF21-Tg) mouse model. Transgenic mice were generated and maintained on a C57BL/6J background. These mice have the FGF21 transgene under the control of apolipoprotein E promoter, which drives the expression of the gene in liver, resulting in overexpression of the FGF21 gene (Inagaki et al., 2007). FGF21-Tg mice have an extended lifespan by about 10 months and have an increased insulin-sensitivity (Zhang et al., 2012). Overexpression of FGF21 increases metabolic rate and induces weight loss in FGF21-Tg mice (Singhal et al., 2016). Moreover, female FGF21-Tg mice are infertile (Zhang et al., 2012).

Since alcohol and saccharin increase dopamine levels in areas of the brain similar to morphine and drugs of abuse, and overexpression of FGF21 has been shown to attenuate preference for alcohol and saccharin, possibly through a dopamine-dependent mechanism (Talukdar et al., 2016), we hypothesized that FGF21-Tg mice would show reduced preference for morphine. Moreover, we also characterized FGF21-Tg and wildtype (WT) mice in assays designed to examine other opioid-induced behavioral and physiological responses including analgesia, tolerance, physical dependence, locomotion, and respiratory depression. Because differences between sexes have been observed in morphine-induced responses in different strains of animals (Hopkins et al., 2004; Kest et al., 1999), we examined both male and female mice in this study.

## 2. Materials and methods

### 2.1. Animals

FGF21-Tg mice and WT littermates (Stock number 021964) were obtained from The Jackson Laboratory, Bar Harbor, ME, USA. Mice were housed five to a cage and maintained in a temperature- and humidity-controlled room at the University of Rochester Medical Center (Rochester, NY) vivarium on a 12:12-h light/dark cycle (lights off at 18:00 h) with food and water available *ad libitum*. Breeding of hemizygous males with WT females yielded approximately 50% FGF21-Tg offspring. Male (234 FGF21-Tg and 266 WT) and female (227 FGF21-Tg and 238 WT) mice (2- to 4-month old) were used for all experiments. Different groups of mice were used for each dose of morphine administration. All procedures were pre-approved and carried out in accordance with the University Committee on Animal Resources at University of Rochester.

### 2.2. Drugs

Morphine sulfate was purchased from Mallinckrodt Chemical Company (St. Louis, MO). Naloxone was purchased from Sigma-Aldrich (St Louis, Missouri, USA). Both morphine and naloxone were solubilized in sterile saline. All injections were administered through intraperitoneal (i.p.) routes in a maximum volume of 10 ml/kg.

### 2.3. Enzyme-linked immunosorbent assay (ELISA) to measure FGF21 protein levels

Serum FGF21 protein levels of FGF21-Tg and WT C57BL/6J littermates were determined using a mouse/rat solid-phase FGF21 Quantikine ELISA Kit MF2100 (R&D Systems, Minneapolis, MN, USA) according to the manufacturer’s protocol. Mouse tail blood was collected in a 1.5-ml Eppendorf tube and left at room temperature for 1 h. The blood was centrifuged at 5000-*g* for 15 min at 25°C. Afterwards, the serum was collected and analyzed by ELISA for FGF21 using an EL800 microplate reader (BioTek Instruments, Winooski, VT, USA) set with a detection wavelength of 450 nm and a correction wavelength at 540 nm. Sample FGF21 concentrations were determined from a standard curve fit with a logistic 4-parameter function using SigmaPlot (Systat Software, Inc.). The mean minimum detectable concentration of FGF21 was 3.81 pg/ml according to the manufacturer.

### 2.4. Conditioned place preference (CPP)

The preference of FGF21-Tg and WT littermates for morphine was determined using a biased CPP protocol as described by Carey et al. (2007). Mice were placed in an apparatus divided into three compartments: two outer compartments (27.3 x 22.2 x 34.9 cm) with distinct wall striping (vertical vs. horizontal alternating black and white lines, 1.5 cm in width) and floor texture (lightly mottled vs. smooth) joined to a smaller central chamber (13.9 x 22.2 x 34.9 cm) with sliding doors (Place Preference, San Diego Instruments, San Diego, CA). Infrared beams along the walls of the apparatus, connected to a computer, allowed automated measurement of time spent in each compartment. Mice were given 30 min to move freely between all compartments to acclimate to the new environment and their initial outer chamber preference was determined. Afterwards, the mice were administered 3 or 10 mg/kg of morphine and place-conditioned in their initially non-preferred chamber for 30 min. Separate groups of mice were treated with different doses of morphine, which were chosen for their reported ability to produce CPP (Hoot et al., 2013; Orsini et al., 2005; Solecki et al., 2009). Six h later, mice were injected with an equivalent volume of saline and placed in their initially preferred chamber for 30 min. This cycle of place conditioning with morphine, followed by saline 6 h later, was repeated the next day, and final place preference was determined on the third day (Hoot et al., 2013). CPP data are presented as the mean difference in time spent in drug- and vehicle-associated chambers ± S.E.M.

### 2.5. 55°C Hot plate and 55°C warm-water tail withdrawal tests to measure antinociception

The antinociceptive responses for FGF21-Tg and WT littermates were compared using ED_50_ values in the 55°C hot plate and the 55°C warm-water tail withdrawal assays. Separate groups of mice were used for each experiment. On the day of the experiment, baseline measurements (basal latencies) were performed before each mouse was administered morphine. The 55°C hot plate test was used to determine the effect of morphine on supraspinal antinociceptive response in FGF21-Tg and WT littermates (Deuis et al., 2017). This test was performed by placing the mouse on a heated surface (55°C, Hot Plate Analgesia Meter, Model 39, San Diego Instruments, San Diego, CA) and measuring the latency for the mouse to jump or lick their paw in response to the heat stimulus. Test latencies were determined at 20, 30, 60, and 90 min after a single morphine injection (18, 20, or 23 mg/kg). A maximum antinociception score (100%) was assigned to mice that did not respond within 30 s. Thus, percentage of antinociception = 100 × [(test latency – basal latency) / (30 – basal latency)].

The effect of morphine on the spinal antinociceptive response in FGF21-Tg and WT littermates was determined using the 55°C warm-water tail withdrawal test (Drgonova et al., 2010; Mathews et al., 2008). Mice were placed in a Plexiglas mouse restrainer and positioned so the tail was immersed in a 55°C warm-water bath. Latency was recorded as the amount of time from when the mouse’s tail enters the water bath to when the mouse perceives a painful stimulus, flicking its tail out of the water (Mathews et al., 2008). Test latency was determined at different time points (20, 30, 60, and 90 min) after morphine administration of 3, 5.6, and 10 mg/kg. A maximum antinociception score (100%) was assigned to mice that did not respond within 15 s to avoid tissue damage (Bidlack et al., 2002; McLaughlin et al., 2006). Percentage of antinociception = 100 × [(test latency – basal latency) / (15 – basal latency)].

### 2.6. Acute morphine analgesic tolerance development

After determination of morphine-induced antinociception in WT and FGF21-Tg littermates, we investigated if FGF21-Tg mice exhibited similar acute morphine-induced analgesic tolerance development as WT littermates using a method described previously (Mathews et al., 2008). FGF21-Tg and WT littermates were treated with 10, 18, or 23 mg/kg morphine in the 55°C hot plate assay and 3, 5.6, or 10 mg/kg morphine in the 55°C warm-water tail withdrawal assay. Morphine was administered at times 0, 3, 6, and 9 h. Time 0 was the first injection of morphine. Dose-response curves showing percentage of antinociception were generated for each time point. Because not every curve reached 50% antinociception, tolerance was determined by comparing the response after time 3, 6 or 9 h to time 0 h at matching doses. Antinociception was assessed 30 min after each injection because morphine produced the greatest antinociceptive response 30 min after injection in the time-course antinociception assays. Percent of antinociception was determined as described above.

### 2.7. Acute morphine physical dependence

To assess the development of acute morphine physical dependence, mice received an injection of a high dose of morphine (100 mg/kg) followed 4 h later by an injection of the opioid antagonist, naloxone (10 mg/kg). This method has been shown to induce development of morphine dependence and withdrawal symptoms such as jumping, forepaw tremor, and rearing (Bilsky et al., 1996; Mathews et al., 2008; Parker and Joshi, 1998; Wang et al., 1999; Yano and Takemori, 1977). We focused on the number of vertical jumps, as this withdrawal symptom was the most apparent. Ten min before injection of naloxone, mice were placed in a large transparent beaker (Height: 125 mm, outside diameter: 35 mm, Kimax Glassware) to habituate them to the new environment. Immediately after the naloxone injection, mice were returned to the beaker and the number of vertical jumps was recorded for 15 min.

### 2.8. Chronic morphine analgesic tolerance development and dependence

In addition to acute morphine analgesic tolerance, we also explored chronic morphine analgesic tolerance development in WT and FGF21-Tg littermates. Mice were treated twice daily (every 12 h) for 5 consecutive days with 23 mg/kg morphine. On day 6, mice were administered a single dose of 23 mg/kg morphine. Analgesia was assessed every day with the 55°C hot plate assay, immediately before and 30 min after the first morphine injection of the day, and the percentage of antinociception was determined. Day 0 was the first day of morphine administration.

On the last day, withdrawal was precipitated with an injection of naloxone (10 mg/kg, 2 h after the last morphine administration). Immediately after the naloxone injection, each mouse was placed in a large transparent beaker and the number of withdrawal jumps was recorded for 15 min.

### 2.9. Locomotion assay

Morphine induces locomotor activity in mice (Valjent et al., 2010). Therefore, we investigated if morphine-induced locomotion would be affected in FGF21-Tg mice in comparison to WT littermates. Saline was used as a control. Mice were administered 10 mg/kg or 23 mg/kg morphine and were placed immediately in an open-field activity apparatus (PAS-Open Field, San Diego Instruments, San Diego, CA) for 2 h. The apparatus uses an open field Plexiglas chamber equipped with photocell emitters and receptors (Tatem et al., 2014). The distance traveled (cm) was recorded for each mouse (Fusion Software, version 3.9, Omnitech Electronics, Inc., Columbus, Ohio).

### 2.10. Morphine-induced respiratory depression

Morphine and other µ opioid agonists activate the µ opioid receptor (MOR) expressed on the surface of neurons in brainstem respiratory centers (Dahan et al., 2010). Activation of these MOR leads to respiratory depression (Pattinson, 2008). To examine if FGF21-Tg mice had a different respiratory rate after morphine administration in comparison to WT littermates, a whole-body plethysmography suite (SCIREQ, Montreal, Canada) was used. This non-invasive method measures mouse respiration using differential pressure transducers connected through an interface (Emka Technologies, France) to a computer for recording and analysis of respiration parameters (Hill et al., 2018). Mice were placed in a small chamber, supplied with room air via a pump, and allowed to move freely throughout the measurements. Based on pressure changes within the chamber, we were able to analyze breathing rates. Mice were habituated to the plethysmography chambers for 3 days for 30 min. On the experimental day (day 3) after the last habituation period, baseline respiration rates, breaths per min (bpm) were measured over a 20-min period before morphine administration. Subsequently, morphine was administered (18 or 30 mg/kg) in a 10-min window and mice were returned to the plethysmograph chambers. The respiration rate was recorded at 5-min intervals for 90 min. Changes in respiration rate were used to evaluate respiratory depression following morphine injection. Data are reported as percent respiration rate relative to baseline ± S.E.M.

### 2.11. Statistical analysis

FGF21 protein expression levels in mice were compared using a 2-way ANOVA (sex x genotype) by IBM SPSS 28.0.0 software. CPP data for male and female mice were analyzed using separate 2-way repeated measures ANOVA (genotype x dose) followed by Sidak-adjusted multiple comparison post hoc tests. Antinociception data for male and female mice were analyzed separately using ANCOVA with genotype as a fixed factor and dose as a covariate. ED50 values and 95% confidence interval (CI) for morphine-induced antinociceptive dose-response curves were created using each individual data point with Prism 5.0 software (GraphPad Software, La Jolla, California). The ED50 values and 95% Cl were compared among the groups. The antinociception time course data from 55°C hot plate experiments were analyzed by 2-way repeated measures ANOVA (genotype x dose) between male WT and FGF21-Tg mice. The same analysis was performed for female mice. Bonferroni adjusted pairwise comparisons were used. Acute morphine antinociceptive tolerance development data were analyzed using 2-way repeated measures ANOVA (genotype x dose) followed by Bonferroni adjusted pairwise comparisons. Differences in acute withdrawal jumps, an assessment of morphine physical dependence, were compared by 2-way ANOVA (sex x genotype) followed by Bonferroni adjusted pairwise comparison. To assess differences in morphine chronic tolerance development between WT and FGF21-Tg mice, 1-way repeated measures ANOVA was used. A 2-way ANOVA was used to compare effects of sex and genotype on chronic withdrawal jumping. Separate 2-way ANOVA (dose x genotype) tests with Bonferroni post hoc multiple comparisons were used to assess locomotor differences within male and female mice. Separate 2-way repeated measures ANOVA (genotype x dose) were used to investigate differences in respiratory rate between WT and FGF21-Tg mice at different doses of morphine within male and female mice. Greenhouse-Geisser adjustment was used when investigating within-subject effects.

## 3. Results

### 3.1. FGF21 protein level in FGF21-Tg and WT mouse serum

FGF21-Tg mice had a 2,400-fold greater level of serum FGF21 than WT littermates as measured by an ELISA (F(1,49) = 677.4, p < 0.001, Table 1). There was no significant effect of sex (p = 0.936) or interaction between sex and genotype (p = 0.934).

**Table 1.** FGF21 serum protein levels from FGF21-Tg and WT littermates. FGF21 protein levels were determined by a FGF21 ELISA assay as described in Materials and Methods (section 2.3.). Data are the mean FGF21 protein levels ± S.E.M. from 10 male and 10 female FGF21-Tg mice and 15 male and 18 female WT mice. There were no differences in FGF21 serum levels between male and female mice of the same strain.

| Mouse Strain | FGF21 (ng/ml) ± S.E.M. |
| --- | --- |
| Male FGF21-Tg | 640 ± 46 |
| Female FGF21-Tg | 650 ± 44 |
| Male WT | 0.30 ± 0.040 |
| Female WT | 0.23 ± 0.045 |

### 3.2. Morphine CPP in FGF21-Tg and WT littermates

FGF21-Tg and WT mice were evaluated for initial preferences in the CPP assay. A 2-way ANOVA (sex x genotype) indicated no significant main effects of sex (F(1,123) = 0.31, p = 0.58) or genotype (F(1,123) = 0.68, p = 0.41) (Fig. 1). In WT mice, morphine at 1 mg/kg did not produce CPP (Supplemental Fig. 1). Morphine CPP was observed following injection of 3 mg/kg and 10 mg/kg morphine (Fig. 1). No significant main effects of genotype or dose were observed for either males or females. There was a significant interaction of time x genotype x dose for males (F(1,57) = 4.04, p = 0.049) and females (F(1,42) = 4.75, p = 0.035). Pairwise comparisons indicated a significant difference between male WT and FGF21-Tg mice in the 10 mg/kg morphine dose (p = 0.012). Similarly, differences between female WT and FGF21-Tg mice were observed at 3 mg/kg (p < 0.001) and 10 mg/kg (p = 0.04) morphine doses. Notably, female FGF21-Tg mice showed no CPP in response to 3 mg/kg morphine (Fig. 1b). These results show that mice expressing high serum levels of FGF21 had a reduced preference for morphine in comparison to WT littermates.

**Fig. 1.**
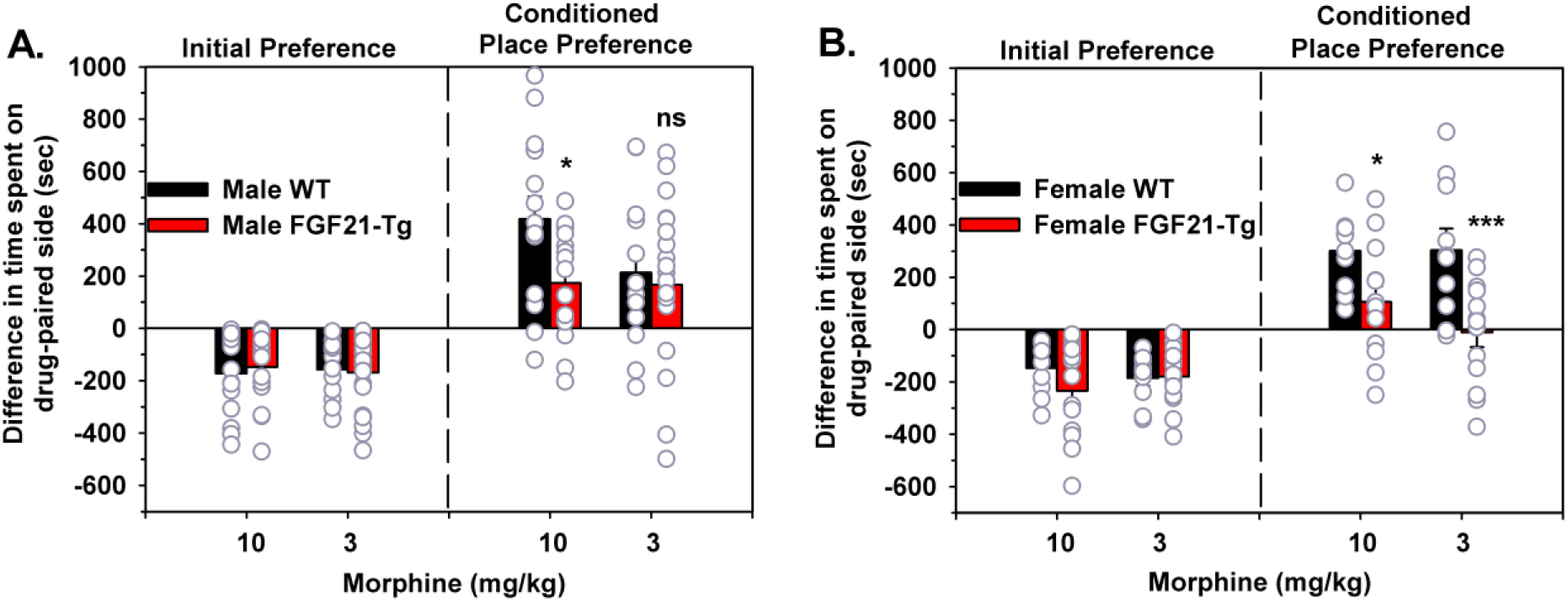
Comparison of FGF21-Tg and WT littermates in morphine CPP. A three-day biased CPP was used in this experiment. Different groups of mice were used for each morphine dose. A) Male FGF21-Tg mice had a lower morphine CPP than male WT mice at 10 mg/kg (*n* = 15-16 mice/group). No significant effect was observed at the 3 mg/kg morphine dose. B) Female FGF21-Tg mice showed reduced preference for morphine at 10 mg/kg and 3 mg/kg morphine in comparison to female WT littermates (*n* = 10-13 mice/group). ns, not significant, *p < 0.05 and ***p < 0.001 for FGF21-Tg mice compared with WT littermates at equal dose.

### 3.3. Morphine-induced antinociception in FGF21-Tg and WT littermates as measured in the 55°C hot-plate and 55°C tail withdrawal assays

Morphine-induced antinociception was measured in FGF21-Tg and WT mice using both the 55°C hot plate and the 55°C warm-water tail withdrawal assays to assess supraspinal and spinal antinociception, respectively. In the 55°C hot plate assay, the mean basal latencies were similar for both FGF21-Tg (7.04 ± 0.08 s) and WT (7.18 ± 0.08 s) mice. 2-Way ANOVA revealed no significant effects of genotype (F(1,116) = 1.6, p =0.21), sex (F(1,116) = 3.1, p = 0.08), nor a significant interaction (F(1,116) = 2.9, p = 0.09). Both FGF21-Tg and WT mice administered 18, 20, or 23 mg/kg morphine reached maximal morphine-induced antinociception 30 min post-injection (Supplemental Fig. 2). Therefore, the 30-min morphine antinociceptive response was used to construct morphine dose-response curves. Morphine produced dose-dependent antinociception. In male mice, a significant effect of dose (F(1,56) = 185, p < 0.001), but not genotype (p = 0.134) or genotype x dose interaction (p = 0.14), was observed, indicating that the slopes of the dose-response curves were not significantly different. The ED_50_ values and 95% CI were 19.4 mg/kg (16.2 - 23.2) and 19.1 mg/kg (15.5 - 23.5) in male FGF21-Tg and WT mice, respectively (Fig. 2a). In female mice, there was a significant effect of dose (F(1,56) = 71.3, p<0.001), but no significant genotype (p = 0.251) or genotype x dose interaction (p = 0.369) were observed. In female mice, the ED_50_ values were 19.6 mg/kg (19.5 - 19.7) for FGF21-Tg and 18.3 mg/kg (16.2 - 20.7) for WT mice (Fig. 2b). Overall, FGF21 overexpression had no effect on supraspinal morphine analgesic response in either male or female mice.

**Fig. 2.**
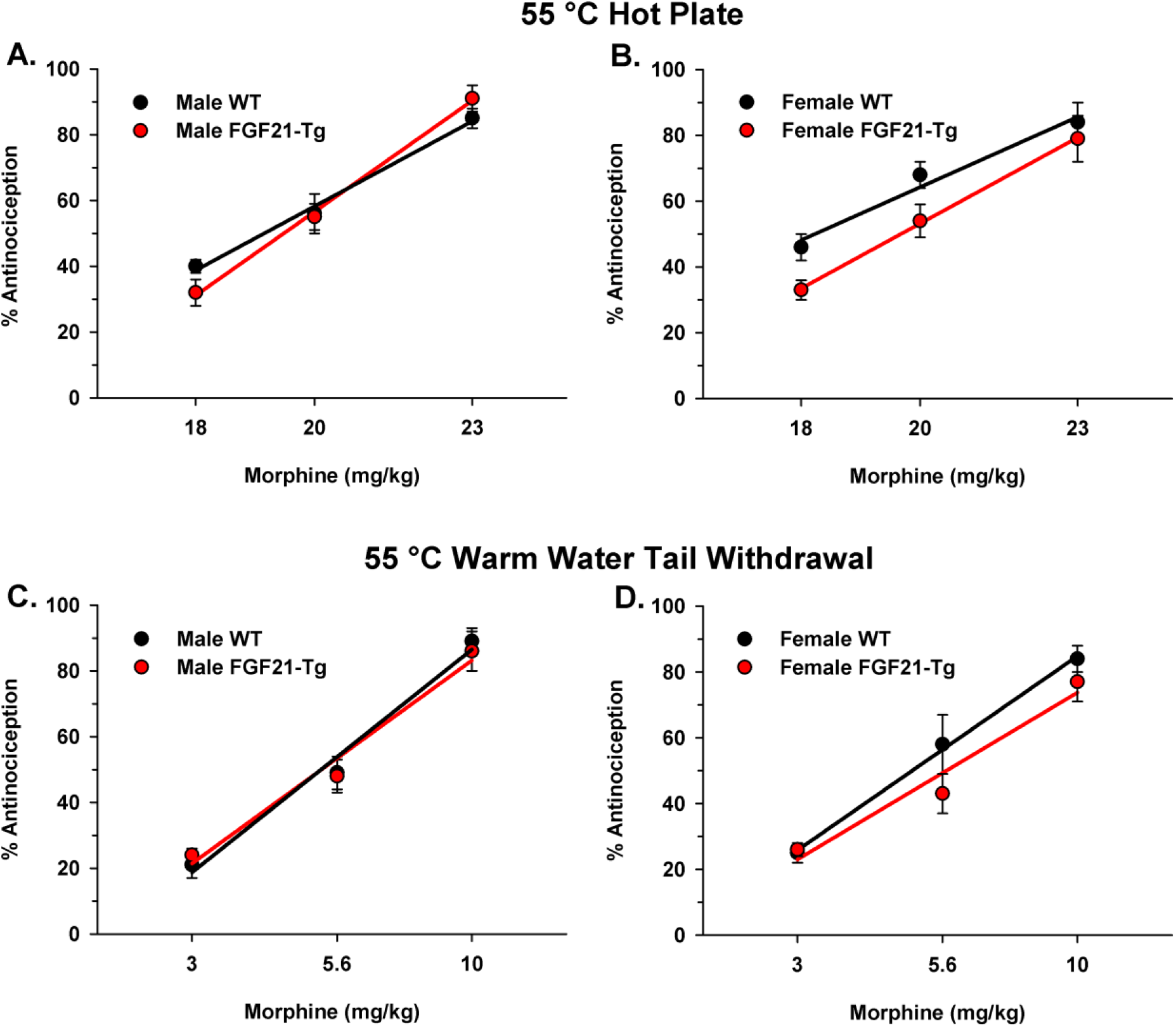
Morphine antinociception in 55°C hot plate (A, B) and 55°C warm-water tail withdrawal (C, D) tests measured 30 min after an i.p. morphine injection. Results (*n* = 10 mice/group) for male (A) and female (B) mice in the 55°C hot-plate test were analyzed separately. Data from the 55°C tail withdrawal test (*n* = 10-15 mice/group) were analyzed separately for male (C) and female (D) mice.

The 55°C warm-water tail withdrawal assay was used to assess whether FGF21 overexpression affects the spinal morphine analgesic response. Mice used in the tail withdrawal assay had similar mean basal latencies of 1.74 ± 0.05 s and 1.70 ± 0.05 s for FGF21-Tg and WT mice, respectively. 2-Way ANOVA indicated no significant effects of genotype (F(1,121) = 0.36, p = 0.55) or sex (F(1,121) = 0.058, p = 0.81). There was a significant interaction (F(1,121) = 4.0, p = 0.05). However, examination of simple effects yielded no significant comparisons. Maximal morphine antinociception was reached 30 min after injection in both transgenic and WT mice at every dose (data not shown). Morphine dose-response curves were constructed similarly to those from the hot plate antinociception assay. In male mice, there was a significant effect of dose (F(1,61) = 193.5, p < 0.001), but no significant effect of genotype (p = 0.654) or genotype x dose interaction (p = 0.571), indicating that the slopes of the dose-response lines were not significantly different from each other. Female mice showed a significant effect of dose (F(1,56) = 71.3, p < 0.001), but no significant effect of genotype (p = 0.840) nor genotype x dose interaction (p = 0.609), indicating the slopes of the dose-response lines were not significantly different. The ED_50_ values and 95% CI were 5.33 mg/kg (1.59 - 17.8) and 5.33 mg/kg (1.85 - 15.4) in male FGF21-Tg and WT mice, respectively (Fig. 2c). These values were 5.85 mg/kg (1.69 - 20.2) and 4.93 mg/kg (4.38 - 5.55) in female FGF21-Tg and WT mice, respectively (Fig. 2d). These data show that there was no difference in the potency or efficacy of morphine in the FGF21-Tg and WT littermates in both antinociceptive assays regardless of sex.

### 3.4. Acute morphine-induced analgesic tolerance development in FGF21-Tg and WT littermates

The development of acute morphine analgesic tolerance in WT and FGF21-Tg mice was measured to determine if FGF21 overexpression had an effect on morphine tolerance development. To evaluate the effect of FGF21 overexpression on supraspinal morphine tolerance development in mice, using the 55°C hot-plate assay, morphine was administered repeatedly at (0, 3, 6, and 9 h) at doses ranging from 18 - 23 mg/kg. Like the antinociceptive assay, mean baseline latencies were measured for FGF21-Tg (7.30 ± 0.07s) mice and WT (6.97 ± 0.06s) littermates. Moreover, basal latencies returned to pre-morphine administration levels before each subsequent injection (data not shown). There was a difference in morphine antinociception in all groups of mice. Significant effects of time (F(3,162) = 71.8, p < 0.001), time x dose (F(6,162) = 4.27, p < 0.001) and time x genotype (F(3,162) = 8.23, p < 0.001) were observed. In the acute tolerance 55°C hot-plate test, male mice became more tolerant to morphine over time and the amount of tolerance development was greater at higher doses of morphine. Overall, male FGF21-Tg mice showed reduced acute tolerance development compared to WT littermates. Post hoc tests revealed that male WT mice developed tolerance at 3 h, following the second morphine injection (p < 0.001, Fig. 3a). Male FGF21-Tg mice began showing signs of tolerance at the 6-h time point, following the third morphine injection (p = 0.002, Fig. 3b). In female WT mice, significant effects of time (F(3,162) = 79.3, p < 0.001) and time x genotype (F(3,162) = 3.41, p = 0.02) and time x dose (FF(6,162) = 4.58, p < 0.001) were observed. Female mice became more tolerant to morphine over time and the amount of tolerance development was greater at higher morphine doses. Overall, female FGF21-Tg mice showed reduced acute tolerance development compared to WT littermates. Female WT mice developed tolerance at 3 h, following the second morphine injections (p < 0.001, Fig. 3c). Female FGF21-Tg mice began showing signs of tolerance at 6 h, following the third morphine injection (p < 0.001, Fig. 3d).

**Fig. 3.**
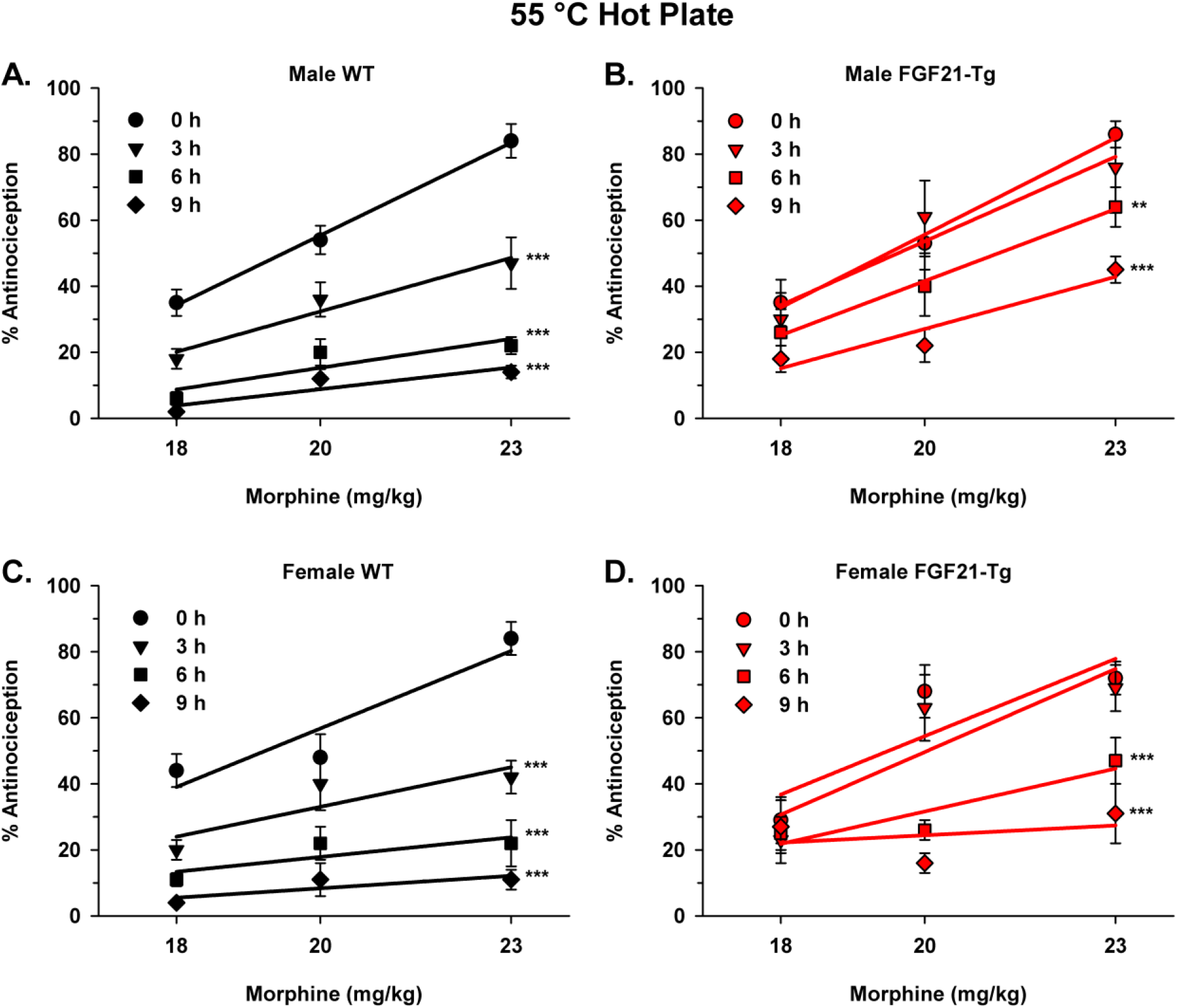
Comparison of FGF21-Tg and WT mice in acute morphine analgesic tolerance development in the 55°C hot-plate test. Mice were administered the same morphine dose (10, 18, or 23 mg/kg) at time 0, 3, 6, and 9 h. Antinociception was assessed 30 min after each injection. A) Male WT mice started to develop tolerance by the second injection (3 h), when administered 23 mg/kg morphine (*n* = 10). B) Male FGF21-Tg mice developed morphine tolerance by the third injection (6 h) when administered 23 mg/kg (*n* = 10). C) Female WT mice became tolerant to morphine by the second injection (3 h) at 23 mg/kg (*n* = 10). D) Female FGF21-Tg mice developed tolerance to morphine by the third injection at 23 mg/kg (*n* = 10). **p < 0.01 and ***p < 0.001 for injection times compared to time 0 h at 23 mg/kg morphine.

A 55°C warm-water tail withdrawal assay was performed to determine the effect of FGF21 overexpression on spinal acute morphine analgesic tolerance development in mice. Morphine was administered by i.p. injection with doses ranging from 3 - 10 mg/kg at (0, 3, 6, 9 h). Baseline latencies for each group of mice used in the tail withdrawal assay were 1.62 ± 0.04 s and 1.61 ± 0.03 s for FGF21-Tg and WT littermates, respectively. There were significant effects of time (F(3,171) = 83.2, p < 0.001), time x dose (F(6,171) = 10.1, p < 0.001), and time x genotype (F(3,171) = 12.2, p < 0.001). Like the hot-plate results, male mice became more tolerant to morphine over time and the amount of tolerance development was greater at higher doses of morphine (Fig. 4a, b). FGF21-Tg mice showed reduced acute tolerance development compared to WT littermates (Fig. 4b). Male WT mice developed tolerance at 3 h, following the second morphine injections (p < 0.001, Fig. 4a). Male FGF21-Tg mice began showing signs of tolerance at the 6-h time point, following the third morphine injections (p = 0.002, Fig. 4b). With female WT and FGF21-Tg mice, significant effects of time (F(3,165) = 79.0, p < 0.001), time x genotype (F(3,165) = 12.9, p < 0.001) and time x dose (F(6,165) = 9.49, p < 0.001) were observed. Like male mice, female mice became more tolerant to morphine over time and the amount of tolerance development was greater at higher morphine doses. Female FGF21-Tg mice showed reduced acute tolerance development compared to WT littermates. Female WT mice developed tolerance at 3 h, following the second morphine injection (p < 0.001, Fig. 4c). Female FGF21-Tg mice began showing signs of tolerance at the 9-h time point, following the fourth morphine injection (p < 0.001, Fig. 4d). The results between the 55°C hot plate and tail withdrawal assays were consistent.

**Fig. 4.**
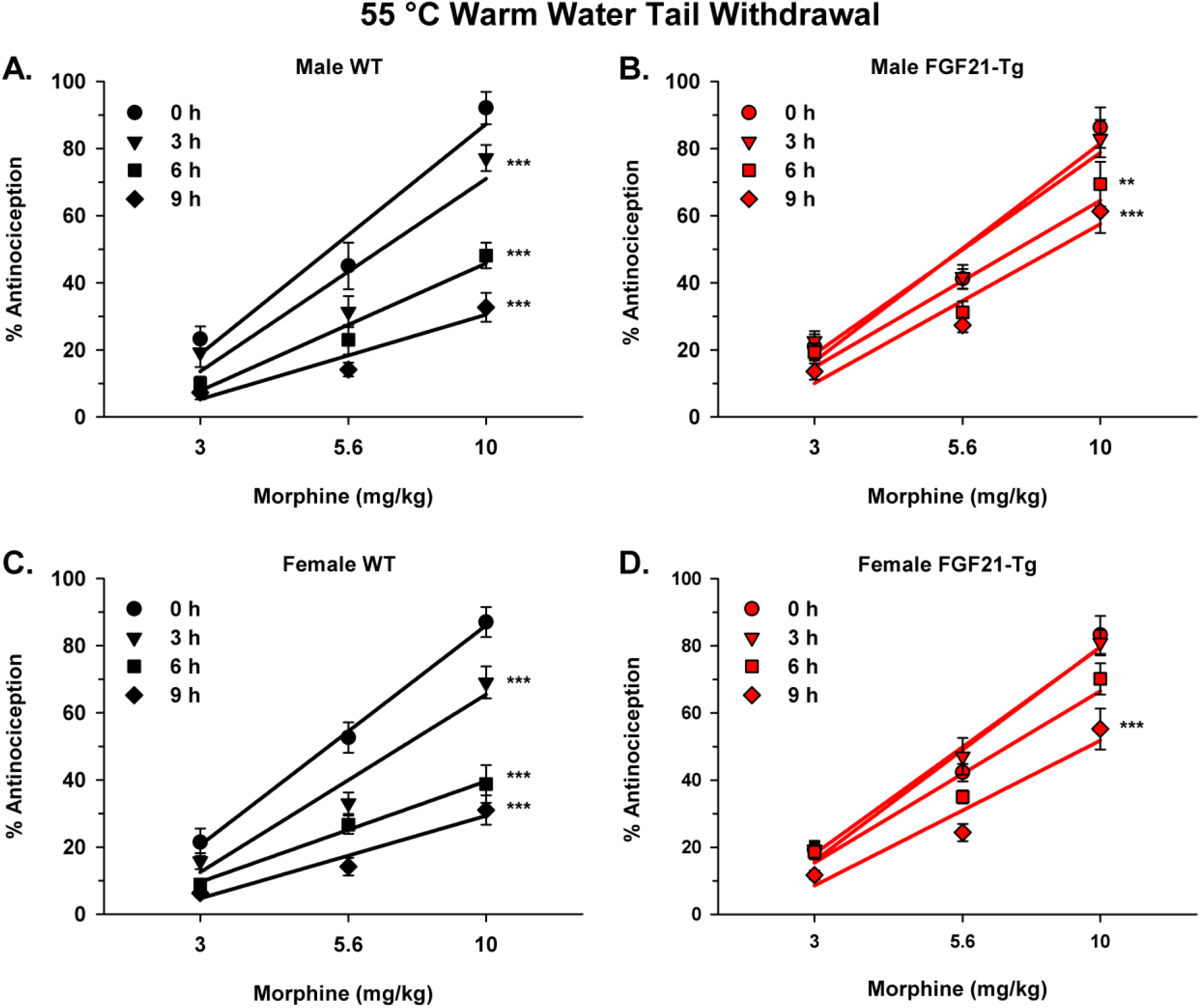
Effect of FGF21 overexpression on acute morphine analgesic tolerance development in the 55°C warm-water tail withdrawal test. Mice were administered the same morphine dose (3, 5.6, or 10 mg/kg) at time 0, 3, 6, and 9 h. Antinociception was assessed 30 min after each injection. A) Male WT mice started to develop tolerance by the second injection when administered with 10 mg/kg morphine (*n* = 10). B) Male FGF21-Tg mice developed morphine tolerance only after the third and final injection of 10 mg/kg morphine (*n* = 10). C) Female WT mice became tolerant to morphine by the second injection (3 h) at 10 mg/kg (*n* = 10). D) Female FGF21-Tg mice showed tolerance after the last injection at 10 mg/kg (*n* = 10). **p < 0.01 and ***p < 0.001 for injection times compared to time 0 h at 10 mg/kg morphine.

### 3.5. Comparison of FGF21-Tg and WT littermates in the development of acute morphine dependence

To understand the effect of FGF21 overexpression on morphine dependence development, an acute morphine dependence paradigm was used (Mathews et al., 2008). The number of withdrawal jumps of morphine-treated mice subsequently injected with naloxone or saline was determined (Fig. 5). Mice treated with saline did not exhibit withdrawal jumping. Acute morphine dependence was observed in all mice, and there was a significant difference based on sex (F(1,36) = 9.91, p = 0.003) and genotype (F(1,36) = 157, p < 0.001) and no significant interaction of these two factors. FGF21-Tg mice displayed fewer withdrawal jumps relative to WT mice. Female FGF21-Tg mice jumped more than male FGF21-Tg mice, following naloxone administration (p = 0.005). No difference was observed between male and female WT mice (p = 0.15).

**Fig. 5.**
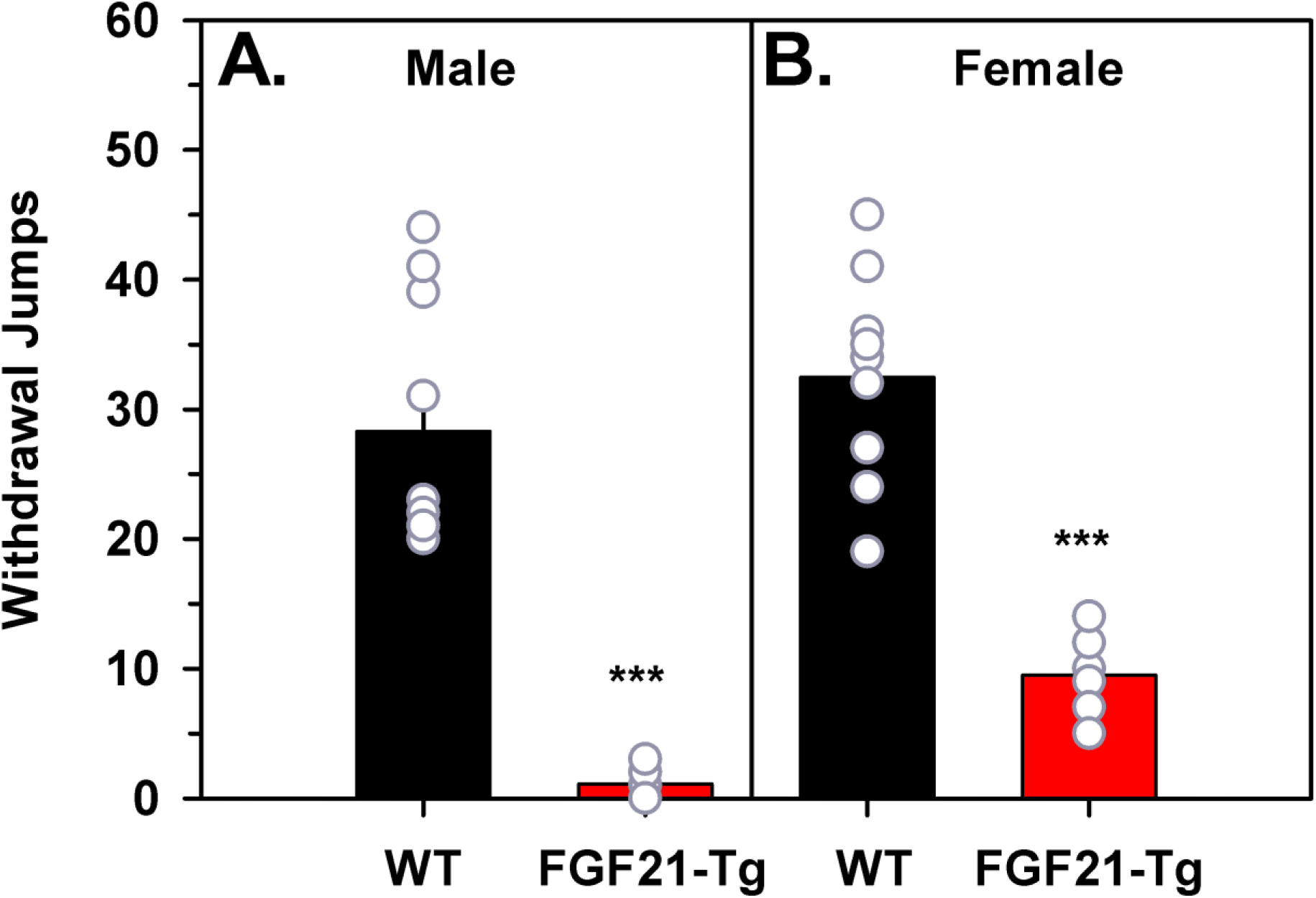
Acute morphine dependence measured in FGF21-Tg and WT mice. Mice received an injection of morphine (100 mg/kg) followed by naloxone (10 mg/kg) 4 h later. Subsequently, mice were observed for 15 min and naloxone-induced withdrawal jumps were recorded. A) Male FGF21-Tg mice had lower withdrawal jumps than male WT mice (*n* = 10 mice/group). B) Female FGF21-Tg had fewer withdrawal jumps relative to female WT mice (*n* = 10 mice/group). ***p < 0.001 for FGF21-Tg mice compared with WT littermates.

### 3.6. Chronic morphine-induced antinociceptive tolerance development in FGF21-Tg and WT littermates

Following the acute morphine-induced analgesic tolerance, we sought to evaluate if FGF21 had any effect on chronic morphine tolerance development (Fig. 6). Mice were administered a dose of 23 mg/kg morphine twice a day for five consecutive days. On the sixth day, mice were administered a single 23 mg/kg dose of morphine. The 55°C hot plate was used to measure antinociception 30 min before and after the first daily morphine dose. FGF21-Tg mice and WT littermates had baseline latencies on day 0 of 7.40 ± 0.17 s and 7.42 ± 0.25 s, respectively. The response latencies after morphine administration decreased in all mice over the 6-day period, demonstrating the development of morphine tolerance after long-term morphine exposure. For male mice, a significant time effect (F(5,75) = 115, p < 0.001) was observed, but there was no significant interaction between time and genotype (F(5,75) = 0.196, p = 0.963), indicating that morphine antinociceptive tolerance developed in both groups of male mice, but there was no significant difference in the responses of WT and FGF21-Tg mice (Fig. 6a). For female mice, a significant time effect (F(5,80) = 97.9, p < 0.001) was observed. However, there was no significant interaction between time and genotype (F(5,80) = 0.552, p = 0.621), indicating that morphine antinociceptive tolerance developed in both female groups of mice, but there was no difference in the responses of WT and FGF21-Tg mice. The development of chronic morphine tolerance was independent of elevated FGF21 levels.

**Fig. 6.**
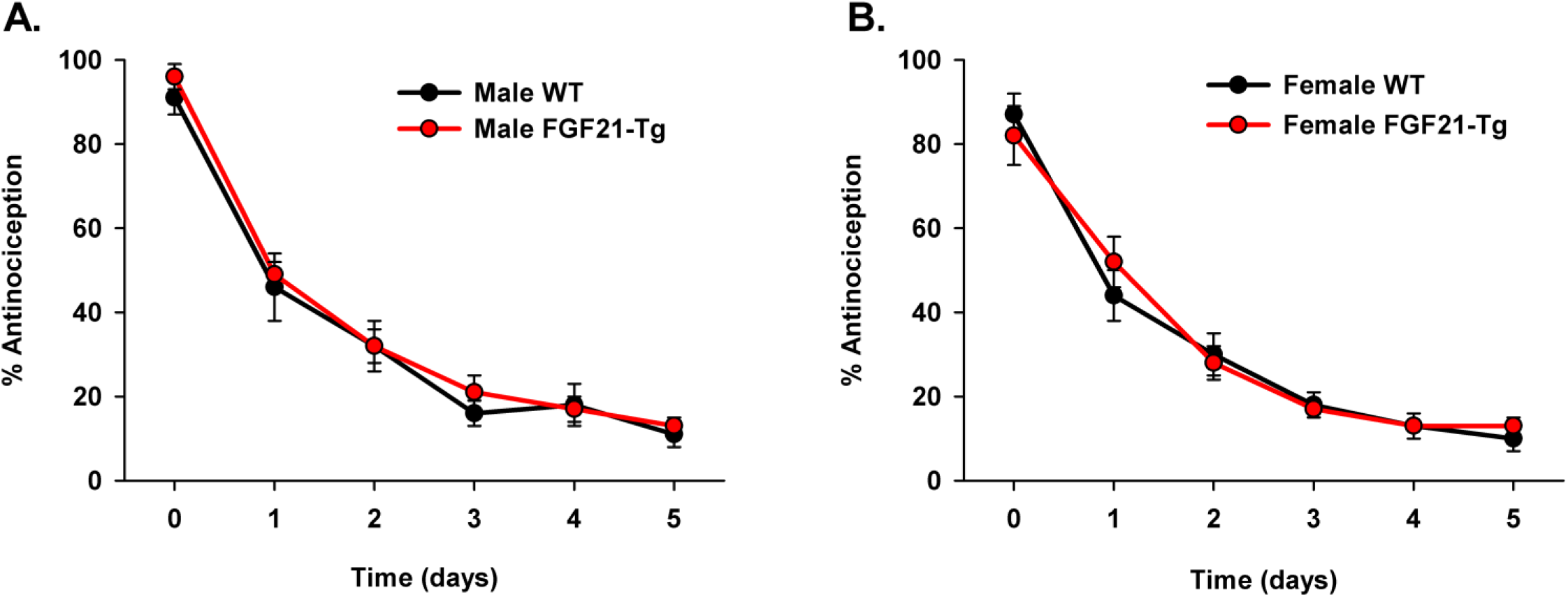
Effect of FGF21 overexpression on chronic morphine analgesic tolerance development in the 55°C hot-plate test. Mice (*n* = 7-10 mice/group) were treated twice a day for 5 consecutive days with 23 mg/kg morphine. Day 0 was the first day of morphine administration. Analgesia was assessed every day, immediately before and 30 min after the first morphine injection. On day 6, morphine-induced analgesia was measured after a single administration of 23 mg/kg morphine. Both male (A) and female (B) WT and FGF21-Tg mice started to develop tolerance by the second day.

### 3.7. Development of chronic morphine dependence in FGF21-Tg and WT littermates

Chronic morphine dependence was evaluated after the last morphine administration of 23 mg/kg on day 5 of the morphine tolerance development model. Mice were treated with naloxone (10 mg/kg) 2 h later, and the number of naloxone-induced withdrawal jumps were observed for 15 min (Fig. 7). A significant main effect of genotype (F(1,31) = 45.5, p < 0.001) was detected, but no significant effects of sex or the interaction of genotype and sex were noted. FGF21-Tg mice displayed fewer withdrawal jumps relative to WT littermates following the development of chronic morphine dependence. Male FGF21-Tg mice withdrawal jumps were reduced by 71% in comparison to male WT littermates (Fig.7a). Similarly, female FGF21-Tg mice withdrawal jumps were 62% lower than female WT littermates (Fig. 7b).

**Fig. 7.**
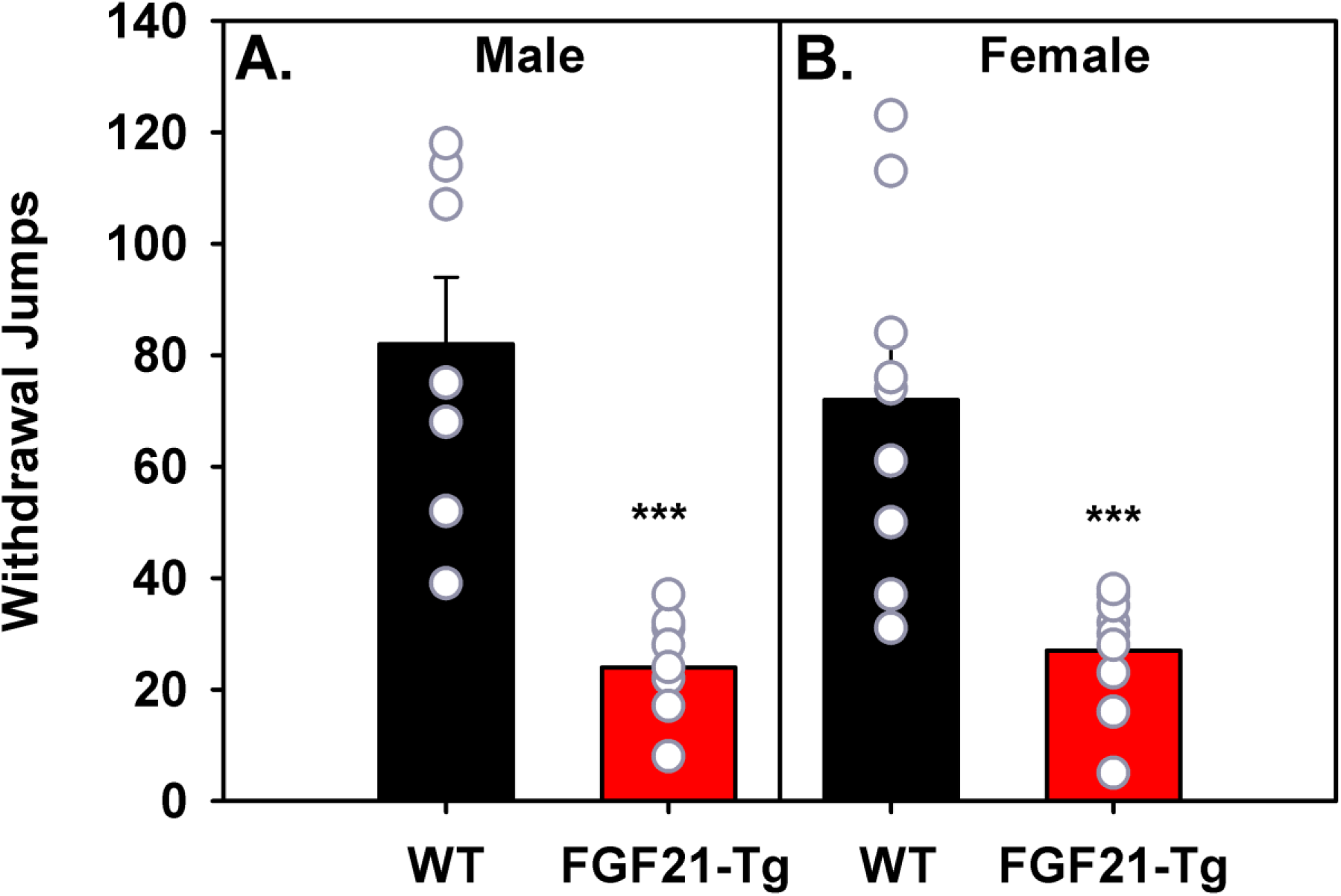
Comparison of male (A) and female (B) FGF21-Tg and WT mice in the development of chronic morphine dependence. Following the development of chronic tolerance on day 5, mice were administered with naloxone (10 mg/kg, 2 h after the last administration of morphine) and observed for 15 min. (*n* = 7-10 mice/group. ***p < 0.001 for FGF21-Tg mice compared with WT littermates.

### 3.8. Morphine-induced locomotion in FGF21-Tg and WT littermates

Morphine induces increased locomotor activity in mice (Hoot et al., 2013; Valjent et al., 2010). The morphine-induced locomotion (distance traveled, cm) between FGF21-Tg and WT littermates was compared (Fig. 8). FGF21-Tg and WT mice had baseline locomotor activity travelling an average distance of 258 ± 13 cm and 256 ± 17 cm over 2 h, respectively. Saline treatment of mice did not alter locomotor activity (data not shown). In male mice, there were no significant main effects of dose (F(1,36) = 1.24, p = 0.273) or genotype (F(1,36) = 3.35, p = 0.076), nor significant interaction of dose and genotype (F(1,36) = 0.301, p = 0.587). Male WT and FGF21-Tg mice were not significantly different in distance traveled at either 10 mg/kg or 23 mg/kg dose of morphine (Fig. 8a). Female mice showed significant main effects of dose (F(1,36) = 15.9, p < 0.001) and genotype (F(1,36) = 35.0, p < 0.001). The interaction between dose and genotype was not significant (p = 0.11). FGF21-Tg female mice showed reduced locomotor performance, in terms of distance traveled, relative to WT female mice. Pairwise comparisons with Bonferroni adjustment indicated that female FGF21-Tg mice had enhanced locomotor activity at the 23 mg/kg morphine dose relative to the 10 mg/kg dose (p < 0.001, Fig. 8b). Distance traveled by WT female mice was not significantly different between the 10 mg/kg and 23 mg/kg morphine doses (p = 0.11).

**Fig. 8.**
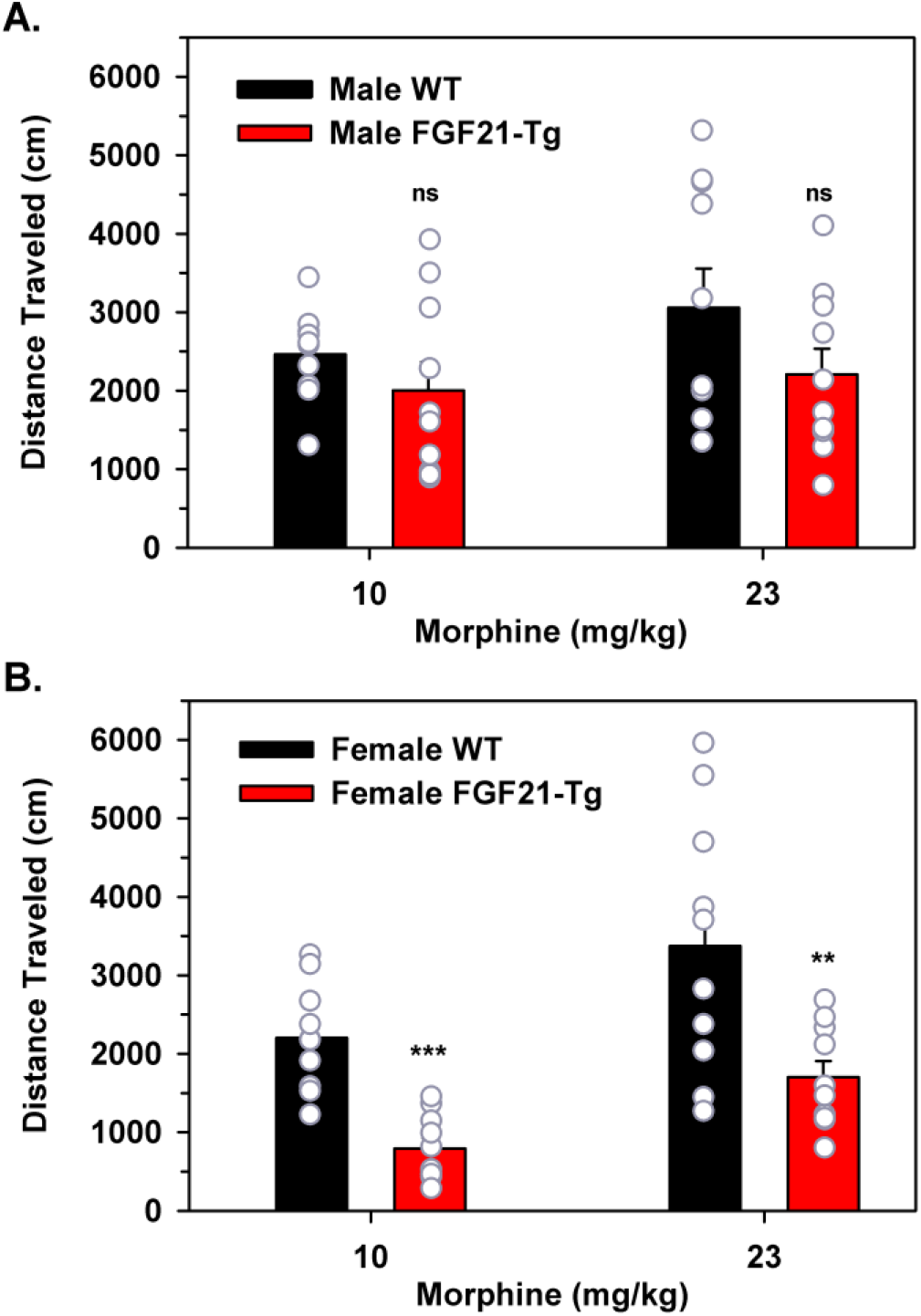
Effects of FGF21 overexpression on morphine-induced locomotion. Male (A) and female (B) FGF21-Tg and WT littermates (*n* = 10 mice/group) were administered with 10 mg/kg or 23 mg/kg morphine and placed in an open-field activity apparatus for 2 h. ns, not significant, **p < 0.01 and ***p < 0.001 for FGF21-Tg mice compared with WT littermates.

### 3.9. Comparing FGF21-Tg and WT littermates on morphine-induced respiratory depression

Morphine produces dose-dependent respiratory depression rapidly after drug injection (Hill et al., 2018). The effect of morphine (18 and 30 mg/kg) on mouse respiratory rate was compared in FGF21-Tg and WT mice (Fig. 9). In male mice, a significant effect of time (F(3, 60) = 54.8, p < 0.001), and a significant time x dose effect (F(3.5, 60) = 2.68, p = 0.05), but no effect of time x genotype (p = 0.16) nor time x dose x genotype interaction (p = 0.812). In male mice, a greater morphine dose resulted in greater respiratory depression, but there was no difference in respiratory response to morphine between male WT and FGF21-Tg mice (Fig. 9a). The respiratory rate did not change after 20 min and persisted over the 90-min observation in both FGF21-Tg and WT mice. Respiration started to recover by 80 min in the mice that received 18 mg/kg, indicative of morphine’s metabolism. In female mice, the results were similar (Fig. 9b). Time was significant (F(3.5,73) = 61.9, p < 0.001), and time x dose was significant (F(3.8, 73.4) = 3.22, p = 0.018), indicating that female mice exhibited a reduced respiratory rate in response to morphine that was exacerbated at the higher morphine dose. The decrease in respiratory rate was independent of the FGF21 protein levels.

**Fig. 9.**
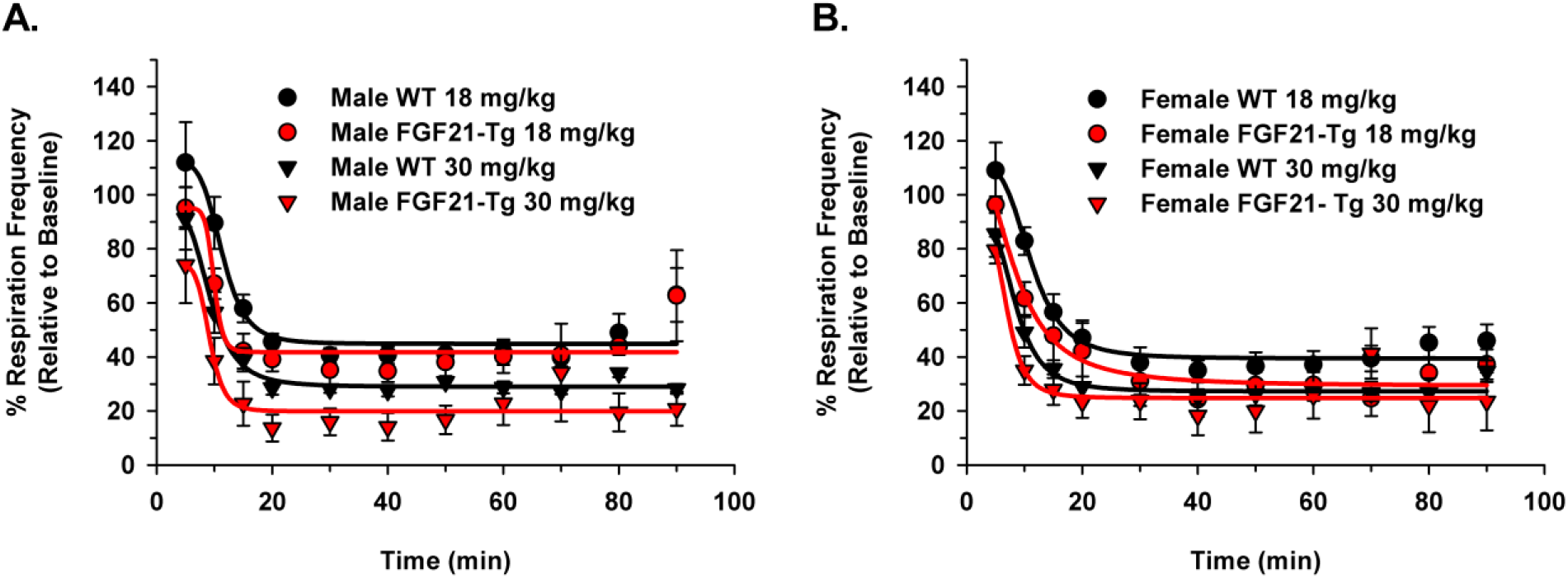
Morphine-induced respiratory depression in male (A) and female (B) WT and FGF21-Tg mice. Mice (*n* = 5-7 mice/group) were placed in a chamber of a whole-body plethysmography suite before and after administration of morphine. Mice breathing rates were analyzed based on pressure changes within the chamber. There were no significant differences between males and female mice and FGF21-Tg and WT littermates.

## 4. Discussion

The goal of this study was to determine if there were any difference in morphine-induced physiological and behavioral responses in FGF21-Tg mice and WT C57BL/6J littermates. We studied morphine-induced events including CPP, acute and chronic tolerance development and dependence, locomotion, and respiratory depression in both male and female mice.

Female FGF21-Tg mice had reduced preference for morphine at 3 and 10 mg/kg doses. However, male FGF21-Tg mice had a lower preference for morphine only at the 10 mg/kg dose. The rewarding effects of morphine are regulated by the CNS, specifically the ventral tegmental area and the NAc (Wilson-Poe and Morón, 2018). Morphine reinforcing effects have been linked to an increase in extracellular dopamine levels in the NAc shells (Fadda et al., 2003). However, Talukdar et al. (2016) reported that mice treated by an osmotic mini-pump containing FGF21 for two weeks had a decrease in dopamine and its metabolites in the NAc relative to vehicle-treated mice. Additionally, FGF21 administration in mice affected the expression of dopamine-related genes, including an increase in the dopamine transporter in the NAc (Talukdar et al., 2016). These findings suggest that FGF21 has a role in the regulation of dopamine in the reward areas of the brain, which correlate with our morphine preference studies. However, additional experiments are essential to confirm if FGF21 regulates morphine-induced dopamine release and to establish the mechanism of action.

FGF21-Tg mice showed no significant difference in basal nociception and in morphine spinal and supraspinal antinociception compared to WT mice. While we did not observe any sex differences in morphine-induced antinociception, 10 mg/kg of oxycodone produced a greater antinociceptive effect in a 54 °C hot plate assay in male C57BL/6J mice than female mice (Collins et al., 2016). The MOR has been shown to regulate the spinal and supraspinal analgesic response of opioids such as morphine (Wang et al., 1994) and mice lacking the MOR did not exhibit morphine-induced antinociception in either the hot plate or the tail withdrawal assays (Kieffer, 1999). FGF21-Tg mice exhibiting similar morphine antinociception compared to WT littermates suggest that the overexpression of FGF21 has no direct effect on the MOR signaling in the spinal and supraspinal regions mediating analgesia.

FGF21-Tg mice displayed reduced development of acute morphine tolerance but similar chronic morphine tolerance compared to WT littermates. FGF21 may modulate acute morphine tolerance development and findings from the following studies may suggest possible mechanisms. Inflammatory cytokines and microglial activation have been linked to morphine tolerance (Wang et al., 2009). Furthermore, blocking interleukin-1 receptor or inhibiting interleukin 1β (IL-1β), a major pro-inflammatory cytokine, was effective in blocking the development of morphine tolerance (Chen et al., 2012). Recombinant human FGF21 suppressed the expression levels of major pro-inflammatory cytokines such as IL-1β, IL-6, and tumor necrosis factor-α (Wang et al., 2020). FGF21 also inhibited microglial activation in mouse hippocampus and the signaling pathway of nuclear factor-κB (NF-κB), another regulator of inflammatory cytokines (Wang et al., 2020). Inhibitors of nitric oxide synthase reduced the development but not the magnitude of acute morphine antinociceptive tolerance (Wolińska et al., 2021; Xu et al., 1998). Under certain circumstances, such as ischemia, FGF21 activated endothelial nitric oxide synthase (Kawanishi et al., 2020). FGF21 effects on acute morphine tolerance may be occurring through some of these pathways.

Phospholipid systems have been associated with opioid tolerance development. Protein kinase C (PKC) inhibitors, 1-(5-isoquinolinesulfonyl)-2-methylpiperazine (H7) and calphostin C, prevented or reversed acute antinociceptive tolerance (Bilsky et al., 1996; Narita et al., 1995). Moreover, PKC knockout mice displayed a decreased morphine tolerance (Zeitz et al., 2001). Mice on a high-fat diet treated with FGF21 via mini-osmotic pumps had reduced PKCϵ activation in liver and PKCθ in skeletal muscle (Camporez et al., 2013). Likewise, FGF21 inhibited the activation of stress-related kinases including NF-κB and PKC in human skeletal muscle (Lee et al., 2012). Although we do not know if the effects of FGF21 on PKC in skeletal muscle and liver also occur in the CNS, FGF21 effects on PKC are an important finding and another potential mechanism by which FGF21 overexpression may affect acute morphine tolerance development. Based on the attenuation of acute morphine tolerance development in mice expressing high FGF21 levels, analogues of FGF21 may be useful in treating acute pain patients receiving opioids to minimize tolerance development.

Chronic morphine tolerance differs from acute tolerance with regards to the length of morphine exposure and possibly the differences in the mechanisms of intracellular modifications of opioid receptors and their second messengers. Chronic activation of the MOR leads to desensitization and downregulation of the MOR. If FGF21 affects acute morphine tolerance through second messenger systems, these effects are conceivably subdued over time and no longer significant during chronic exposure to morphine. Thus, the changes in the number of MOR and signaling efficacy during chronic stimulation by an agonist would not be affected by FGF21 overexpression, explaining the similarity in chronic morphine tolerance development of FGF21-Tg and WT littermates.

FGF21-Tg mice displayed reduced naloxone-precipitated morphine withdrawal symptoms compared to WT mice after being exposed to morphine both acutely and chronically. Acute morphine exposure induces changes in gene expression through decreased expression of coding transcription factors such as c-fos, c-jun, and zif268, which persists with chronic opioid administration (Nestler, 1992). Moreover, Hayward et al. (1990), reported that expression of c-fos is increased during naltrexone-induced opioid withdrawal. These results suggest that c-fos and related transcription factors may regulate neuronal plasticity during morphine exposure. FGF21 has been shown to increase the expression of c-fos in regions associated with opioid dependence (von Holstein-Rathlou et al., 2016). The fact that FGF21-Tg mice had reduced morphine withdrawal symptoms compared to WT mice after being exposed to morphine both acutely and chronically suggests that FGF21 regulation of transcription factors may be involved in morphine physical dependence pathways. FGF21 analogues may help opioid-dependent patients withdraw from opioids by having attenuated withdrawal symptoms. Similarly, craving for opioids may be reduced with FGF21 analogues.

Locomotor activity in response to morphine varied with sex. Female FGF21-Tg mice had reduced morphine-induced locomotor activity compared to WT littermates, but male FGF21-Tg mice had similar morphine-induced locomotor activity to their WT counterparts. In addition, 10 mg/kg morphine produced the maximal effect on locomotor activity in male FGF21-Tg mice but not in female FGF21-Tg mice. The CNS plays an important role in mediating locomotion (Frigon, 2017). The precise site of action of FGF21 in the CNS is unknown. However, its co-receptor, KLB, is expressed in areas such as the suprachiasmatic nucleus, cortex, striatum, hippocampus, and hypothalamus (Lein et al., 2007). Thus, overexpression of FGF21 affects many pathways and hormones, which in turn leads to different phenotypes and genotypes in male and female mice. For example, FGF21-Tg female mice are infertile, but FGF21-Tg male mice are fertile. This phenomenon is the result of FGF21 suppressing vasopressin and kisspeptin signaling (Owen et al., 2013). The difference in morphine-induced locomotion in FGF21-Tg male and female mice may be the result of FGF21 effects on hormonal responses upstream of the locomotion pathway.

It is intriguing that mice with elevated FGF21 levels had the same morphine-induced antinociception and respiratory depression as WT mice. These findings demonstrate that FGF21 does not alter all opioid-induced behavioral responses. From the results of this study, FGF21 and FGF21 analogues may be potential therapeutics for acute pain patients to reduce the development of physical and psychological dependence. Unfortunately, FGF21 has a short half-life (< 2 h) across multiple species limiting its utility (Yie et al., 2009). FGF21 analogues are an excellent way to study the effect of FGF21. PF-05231023 and Efruxifermin are two FGF21 analogues currently in clinical trials (Kaufman, 2020; Lee et al., 2016; Xu et al., 2009). PF-05231023 was developed by covalent conjugation of two molecules of modified FGF21 to a CovX-200 antibody scaffold resulting in two free FGF21 C- and N-termini, respectively (Huang et al., 2013). Both C- and N-termini are required for the formation of the FGF21/KLB/FGFR complex (Kharitonenkov and Larsen, 2011). FGF21 was conjugated to CovX body at the mid-region of the FGF21 protein (Foltz et al., 2012) and the conjugation site has no significant impact on the *in vitro* potency of the protein (Xu et al., 2009). PF-05231023 has a 22 h half-life (Weng et al., 2015). Efruxifermin, formerly AKR-001, Fc-FGF21(RGE), and AMG 876, is a human immunoglobulin Fc-FGF21 fusion protein with a half-life of 3 - 3.5 days (Kaufman, 2020). These and additional FGF21 analogs are being pursued in clinical trials for treating obesity, type 2 diabetes mellitus, and nonalcoholic steatohepatitis (Geng et al., 2020; Tillman and Rolph, 2020). Agonistic monoclonal antibodies specific for the FGF21 receptor complex are being studied (Geng et al., 2020).

The present study showed suggests that FGF21 analogues acting centrally may be therapeutics for treating acute pain patients along with an opioid. The FGF21 analogues may attenuate the development of acute tolerance, and physical and psychological dependence without affecting analgesia.

## 5. Conclusions

In summary, we found that mice expressing high FGF21 serum levels had a reduced preference for morphine, less severe morphine withdrawal symptoms, and slower acute antinociceptive tolerance development than WT littermates. High FGF21 levels did not affect morphine-induced antinociception, chronic antinociceptive tolerance, and respiratory depression. Our results suggest FGF21 and its receptor are therapeutic targets for treating opioid-mediated withdrawal symptoms and craving, and acute opioid tolerance development in patients receiving opioids acutely for pain management.

## CRediT authorship contribution statement

**Louben Dorval:** Conceptualization and design of experiments, data curation, Writing - original draft and revision, was primarily responsible for the collection, interpretation, and analysis of data. **Brian I. Knapp:** Conceptualization and design of experiments, primarily responsible for ELISA and tail-withdrawal assays, Writing - interpretation of the data, review & editing. **Olufolake A. Majekodunmi:** Helped with acute morphine dependence and morphine-induced locomotion. **Sophia Eliseeva:** Helped with morphine-induced respiratory depression. **Jean M. Bidlack:** Conceptualization and design of experiments, Writing - interpretation of the data, reviewing and editing of the manuscript. Funding acquisition. All authors contributed to, and have approved, the final manuscript.

## Acknowledgements

We thank Dr. Steve N. Georas for assistance with the respiratory depression experiments.

## Supplemental Data

**Supplemental Figure 1.**
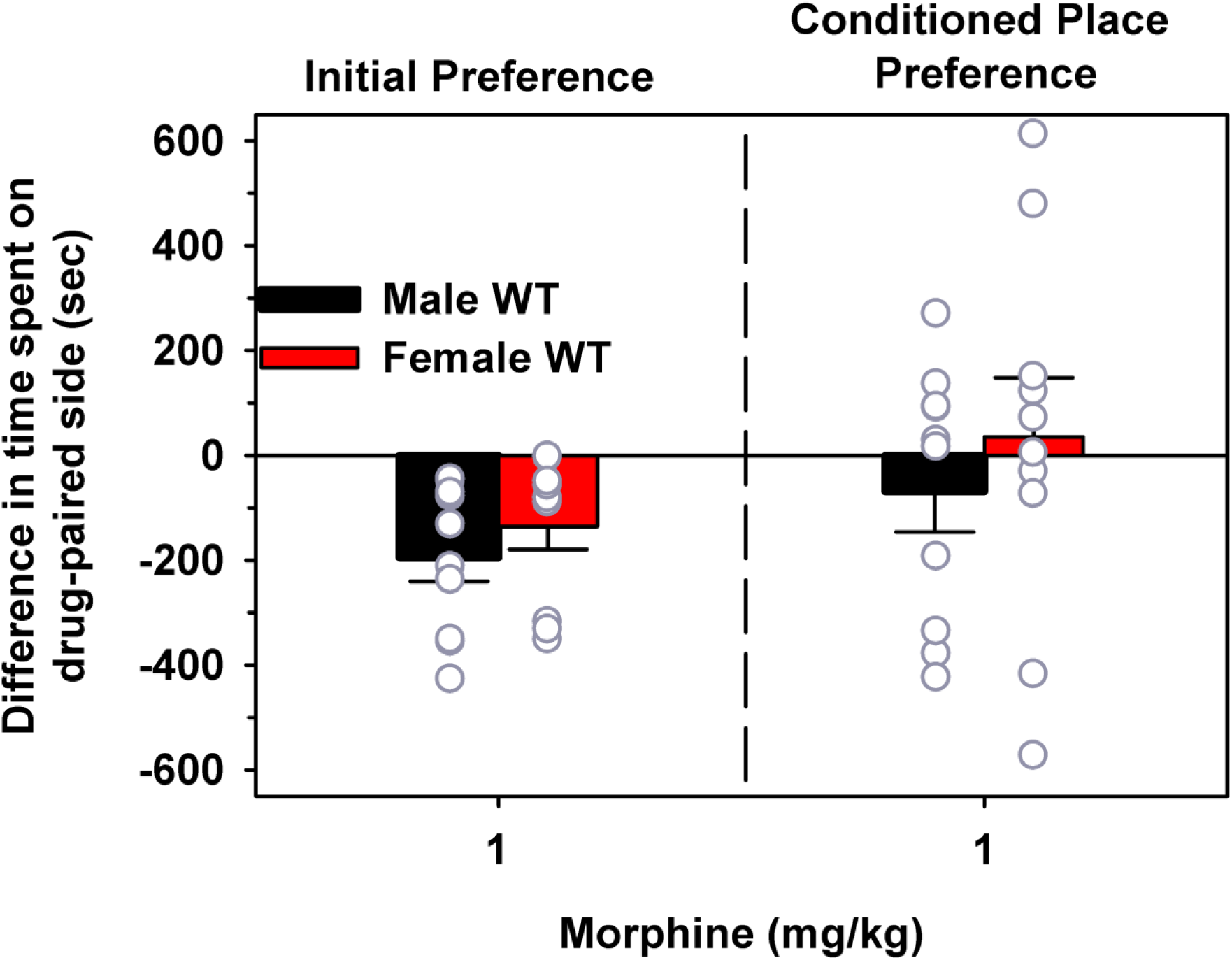
Effect of 1 mg/kg morphine on CPP. A three-day biased CPP was used in this experiment. 1-Way repeated measures ANOVA was used to evaluate differences in initial and conditioned place preference of male and female wild-type mice administered 1 mg/kg i.p. morphine. The analysis indicated no significant time (F(1,18) = 3.26, p = 0.09) effect, nor significant time*sex interaction (F(1,18) = 0.074, p = 0.788) indicating that males and females did not differ in their preconditioning and post-conditioning preference with 1 mg/kg morphine.

**Supplemental Figure 2.**
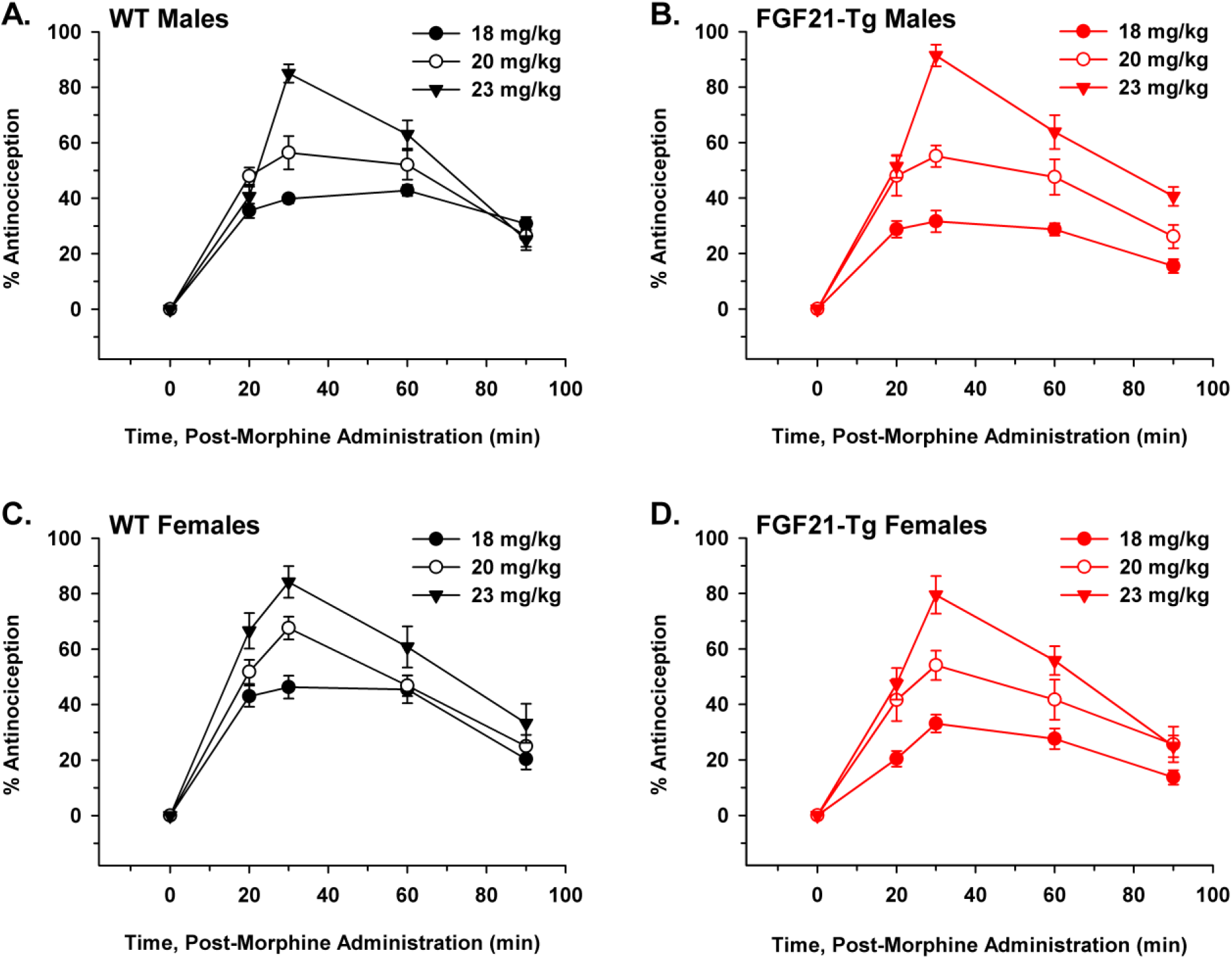
Morphine antinociceptive time course for FGF21-Tg and WT mice in the 55°C hot plate. Morphine produced a peak antinociception 30 min after a single administration in all mice treated with 18-23 mg/kg. 2-Way repeated measures ANOVA (genotype x dose) was used to evaluate differences in 55°C hot plate analgesic responses between male WT and male FGF21-Tg mice, and separately female WT and female FGF21-Tg mice, following i.p. administration of various doses of morphine. The analysis indicated a significant time effect (F(3,159) = 107.8, p < 0.001) and a significant effect of dose x time (F(6,159) = 20.1, p < 0.001), but no effect of genotype x time (F(3,159) = 1.59, p = 0.19) or interaction of time, genotype, and dose (F(6,159) = 0.8, p = 0.574). For female mice, the analysis indicated a significant time effect (F(3,162) = 138.7, p < 0.001), significant time x dose effect (F(6,162) = 8.2, p < 0.001), and significant time x genotype effect (F(3,162) = 4.0, p = 0.009), but no significant time*genotype*dose interaction (F(6,162) = 1.25, p = 0.28). Examination of post hoc pairwise multiple comparisons tests with Bonferroni adjustment indicated that female FGF21-Tg mice had reduced analgesic responses to morphine on average, irrespective of dose, compared to their WT counterparts. These differences were significant at the 20 min (p < 0.001), 30 min (p = 0.013), and 60 min (p = 0.044) time points.

## Abbreviations

ANOVA: analysis of Variance
bpm: breaths per minute
CI: confidence interval
CNS: central nervous system
CPP: conditioned place preference
ED50: median effective dose
ELISA: enzyme-linked immunosorbent assay
FGF: fibroblast growth factor
FGF21-Tg: FGF21 transgenic
H7: 1-(5-isoquinolinesulfonyl)-2-methylpiperazine
IL-1β: interleukin-1β
i.p.: intraperitoneal
KLB: β-Klotho
MOR: mu opioid receptor
NAc: nucleus accumbens
NFκB: nuclear factor kappa B
PKC: protein kinase C
WT: wildtype

